# Effects of phenotypic robustness on adaptive evolutionary dynamics

**DOI:** 10.1101/511691

**Authors:** Emanuele Rigato, Giuseppe Fusco

## Abstract

Theoretical and experimental studies have provided evidence for a positive role of phenotype resistance to genetic mutation in enhancing long-term adaptation to novel environments. With the aim of contributing to an understanding of the origin and evolution of phenotypic robustness to genetic mutations in organismal systems, we adopted a theoretical approach, elaborating on a classical mathematical formalizations of evolutionary dynamics, the *quasispecies model*. We show that a certain level of phenotypic robustness is not only a favourable condition for adaptation to occur, but that it is also a necessary condition for short-term adaptation in most real organismal systems. This appears as a threshold effect, i.e. as a minimum level of phenotypic robustness (critical robustness) below which evolutionary adaptation cannot consistently occur or be maintained, even in the case of sizably selection coefficients and in the absence of any drift effect. These results, are in agreement with the observed pervasiveness of robustness at different levels of biological organization, from molecules to whole organisms.

## 1 Introduction

Phenotypic robustness has been defined as the property of a biological system to preserve its phenotype in the face of perturbations, such as genetic mutations (Wagner, 2011, 2013). This quality is widely considered to be pervasive at different levels of biological organization, from molecules to whole organisms (Kitano, 2004; Stelling et al., 2004; Wagner, 2005). At the level of the organism, phenotypic robustness to genetic mutations might seem to be a quality of the organism’s genotype-phenotype (G-P) map that should hamper the process of adaptation, by making the occurrence of beneficial phenotypic mutations more rare. However, somewhat counter-intuitively, theoretical and computational studies predict a positive role for phenotypic robustness in enhancing long-term adaptation to novel environments (Gibson and Reed, 2008; Wagner, 2008; Draghi et al., 2010; Hayden et al., 2011). This effect has been demonstrated through simulations (Rodrigues and Wagner, 2009; Barve and Wagner, 2013) and experimental studies on ribozymes (Hayden et al., 2011). More recently, an experimental evolution study on *E. coli* has shown that phenotypic robustness can promote significantly faster adaptation at the level of a whole organism (Rigato and Fusco, 2016).

There are two main ways in which phenotypic robustness has considered to be able to foster adaptation (Wagner, 2012) through the accumulation of cryptic genetic variation (CGV). Firstly, in a new environment, phenotypically unexpressed genetic variation can become expressed, and among the new phenotypes, some variants might result accidentally “pre-adapted”, or “exapted” to the new environmental conditions (Hayden et al., 2011). Secondly, and possibly more importantly because less fortuitous, the scattering of genotypes with the same phenotype through the genotype space allows the population to access a greater number of new phenotypes through mutation, increasing the probability of finding phenotypes that happen to have higher fitness (Rodrigues and Wagner, 2009). This latter mode has been recently questioned by Mayer and Hansen (2017), who, on the basis of a computational study based on Boolean networks, argued that positive effects of robustness on evolvability can emerge only under strict biological conditions. However, there is possibly a third way, which is particularly on focus here and takes into account the fact that robustness can support the spread of already present favourable phenotypic variants. As we will show, this is an effect of the further damping of the mutation probability owing to a generic property of the G-P map, that in practice increases the evolutionary stability of phenotypic variants. While the first two aforementioned effects are contingent on the structure of variation of the population, including the level of CGV, and on the structure of the neutral networks in the genotype space, the last one is to a large extent constitutive, i.e. independent on new adaptive challenges (like a new environment, or a modified fitness landscape), and it is able to produce short-term effects.

With the aim of contributing to an understanding of the origin and evolution of phenotypic robustness to genetic mutations in organismal systems, we adopted a theoretical approach, by elaborating on a classical mathematical formalizations of evolutionary dynamics, the *quasispecies model* (Eigen et al., 1989). We derived a phenotypic version of the quasispecies model, which describes frequency dynamics at phenotypic level. Then, by appropriate decomposition, we extracted a phenotypic-robustness term which is significant in the current discussion on the role of robustness in evolution.

## 2 Model assumptions

The quasispecies model is a mutation-selection dynamical system, which describes the evolution of an infinitely large population of haploid, asexually reproducing genotypes on a constant fitness landscape (Nowak, 2006). This is a deterministic model and our derivations are based on three assumptions: i) The view on phenotype is restricted to the *target phenotype*. This is defined as the phenotype that would be expressed by a given genetic makeup of the organism under some given environmental conditions during development, in absence of perturbations (Nijhout and Davidowitz, 2003). This is not to neglect environmental effects on the phenotype, either in the form of phenotypic plasticity or developmental noises (Fusco and Minelli, 2010), but rather to concentrate on the contribution of the organism’s genotype to its phenotype. Thus phenotypic plasticity, i.e. the fact that individual genotypes can produce different phenotypes when exposed to different environmental conditions (Fusco and Minelli, 2010) and developmental instability produced by random perturbations of development are not accounted for here. ii) The genotype includes the whole genome of the organism, as a single allele determining the phenotype (*omnigenic model*; Boyle et al., 2017). This perspective is supported by a double argument. On one side, a phenotypic trait generally depends on the expression of numerous genetic determinants, although with effects of variable magnitude (Fisher’s *infinitesimal model;* Turelli, 2017). On the other side, virtually each locus can, more or less directly, affect a vast array of phenotypic traits (*ubiquitous pleiotropy*; Visscher and Yang, 2016). The omnigenic model is supported by empirical evidence, the most recent coming from genome-wide association studies (GWAS) (review in Boyle et al., 2017), but with reference to our derivations, this choice allows to avoid specific assumptions on more detailed features of the organism’s G-P map, including the level of epistasis, pleiotropy, and neutrality, for which, despite substantial theoretical modelling (Orr, 2000; Wagner, 2008; Wagner and Zhang, 2011; Pavlicev and Wagner, 2012), the few observational data (Grüneberg, 1938; Albert et al., 2008; Rohner et al., 2013) show low levels of generality. iii) A key generic feature of the G-P map at the level of the organism (when the phenotype is intended as target phenotype) is that this is a many-to-one relationship. Stated differently, multiple genotypes can map on the same phenotype (Wagner, 2011; Ahnert, 2017; Mayer and Hansen, 2017).

Elaborating on the quasispecies formalization of evolutionary processes, here we show that a certain level of phenotypic robustness is not only a favourable condition for adaptation to occur, but it is also a necessary (although not sufficient) condition in most real organismal systems. This appears as a threshold effect, i.e. as a minimum level of phenotypic robustness (critical robustness) under which evolutionary adaptation cannot occur or be maintained, even in the case of sizably selection coefficients and in the absence of any drift effects.

## 3 Phenotypic robustness and phenotypic stability

Phenotypic robustness is a property of the genotype-phenotype map. Here, for the derivations to follow, we will adopt a narrow, quantitative definition of *phenotypic robustness* (*ρ*), that is the probability that mutation of a given genotype *g* across one generation, takes to a genotype *g′* that exhibits the same phenotype of *g*.

From this definition of robustness, a definition of *phenotypic stability* (*ϕ_pp_*) follows. This is the probability that the replication of a given genotype *g* with phenotype *p* takes to a genotype that exhibits the same phenotype *p*. Indicating with *η_g_* the mutation probability per genome per generation, phenotypic stability results to be the sum of the probabilities of two mutually exclusive events, namely i) that there is no mutation (1 – *η_g_*) and ii) that in case of mutation the mutant genotype maps to the same phenotype (*ρη_g_*), that is

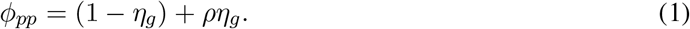

## 4 Quasispecies model analysis

The quasispecies model (Eigen et al., 1989) is a single locus, multi allele, mutation-selection model where each allele differs from the others by at least a single point mutation.

Let’s imagine a sufficiently large population of *n* replicating sequences (or, asexually reproducing haploid genotypes). Sufficiently large means that we can neglect the effects of drift. Sequences can replicate at different rates, according to their fitness and can mutate upon replication. Denote by *x_i_* the relative frequency of the *i*th sequence type, thus we have 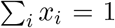. The population structure is given by the vector ***x*** = (*x*_1_, *x*_2_,…*x_n_*). Denote by *q_ji_* the per-replication probability of a sequence *j* to mutate into a sequence *i* and by *W_i_* the absolute fitness (absolute growth rate) of the ith sequence type. The fitness landscape is given by the vector ***W*** = (*W*_1_, *W*_2_,…, *W_n_*) and the average population fitness is 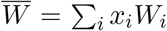 (see Nowak, 2006). In its continuous-time version, the quasispecies equation expresses the time derivative of the frequency of the *i*th sequence type as

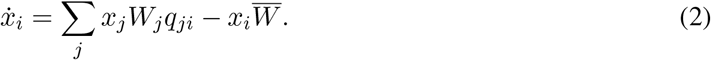

Equation (2) describes the evolution of population of genotypes on an invariant fitness landscape, where the absolute fitness of a genotype does not depend of its own frequency (frequency-independent selection).

### 4.1 Introducing the genotype-phenotype dualism into the quasi species model

Since the principle of the quasispecies dynamics holds for every mutating and reproducing entity, we can use the quasispecies formalism to track frequency changes at phenotypic level, rather than at the level of the genotype. Let’s rewrite the quasispecies equation for a focal phenotype *p*, with frequency *x_p_*, as

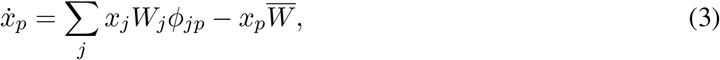

where *W_j_* is the fitness of the *j*th phenotype, *ϕ_jp_* is the phenotypic mutation probability of *j* into *p* and 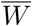 is the population mean fitness 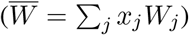. Decomposing the summation in (3) to highlight the two main contributions to the frequency change of *p*, yields

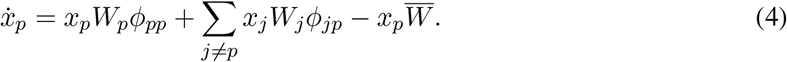

Equation (4) is the phenotypic version of the quasispecies model, assuming different genotypes to map on the same phenotype. The first term of the right-hand side of (4) is the contribution of non-mutant phenotypes *p*, while the second term is the sum of the contribution of mutations from different phenotypes. *ϕ_pp_* is the phenotypic stability term, which in its turn contains the robustness term *ρ*. Derivations similar to equation (3) were developed by Reidys et al. (2001) and Takeuchi et al. (2005). However, having a different aim and moving from different assumptions with respect to the present modellization, these two contributions adopted a quantification of the G-P map redundancy that does not coincide with phenotypic robustness (*ρ*) as defined here (see Discussion).

### 4.2 Conditions for adaptation

Considering equation (4), let’s define that adaptation occurs when an advantageous phenotype *p* (i.e. a phenotype with 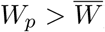) increases its frequency, that is when *ẋ_p_* > 0. Then we can write

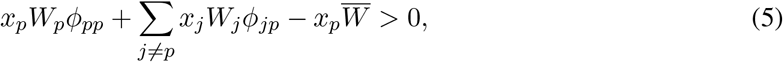

and dividing both therms of the inequality by 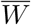, gives

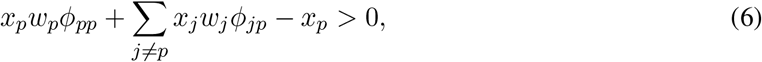

where *w_p_* and *w_j_* indicate the relative fitness of the phenotypes. Note that this definition of adaptation focuses on the instantaneous ability of the population to adapt, and does not require any equilibrium analysis. At variance with most treatments of the quasispecies equation, the advantageous phenotype does not need to be the most advantageous phenotype and the analysis does not assume a closed system (i.e. a system where the arrival of mutant phenotypes that are fitter than the focal phenotype can be ignored).

The mutational contribution from different phenotypes to the frequency of the focal phenotype 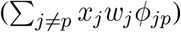 is analogous to the probability of back mutations in the standard application of the quasispecies equation, a term often neglected to simplify further analytical elaborations (e.g., Nilsson and Snoad, 2002; Sasaki and Nowak, 2003; Gorodetsky and Tannenbaum, 2008; Draghi et al., 2011). Here we observe that this frequency value is expected to be even less influential, because of the generally huge size of whole-genome genotype. However, more importantly, this values (always ≥ 0) represents a non-constitutive component of the buffering effect of robustness, being contingent on the specific phenotype, the current distribution of genotypes in the genotype space and other local detailed features of the G-P map. Thus, aiming at concentrating on the constitutive effects of robustness, by setting this term to zero and rearranging we get

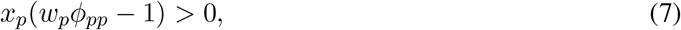

which simplifies to

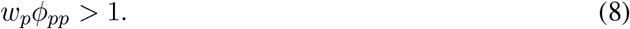

Inequality (8) is the necessary condition to be satisfy for adaptation to occur without relying on unpredictable factors. Since the phenotypic stability term *ϕ_pp_* contains the robustness term, by substituting (1) into (8), the required minimum level of robustness to satisfy (8) results to be

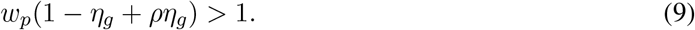

Rewriting the relative fitness term as *w_p_* = (1 + *s_p_*), were *s_p_* is the selection coefficient of the advantageous phenotype *p*, we get

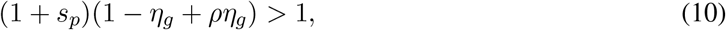

and by isolating the *ρ* term, we finally obtain

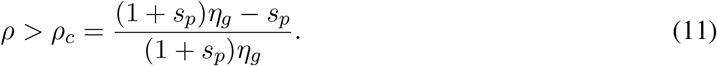

The right-hand side of inequality (11) is the minimum level of phenotypic robustness required for adaptation to consistently occur or to be maintained under the quasispecies model, that we indicates as the *critical robustness* (*ρ_c_*). This depends exclusively on the genome mutation probability *η_g_* and on the selection coefficient *s_p_*. As the mutation probability increases, higher levels of phenotypic robustness are required for adaptation to occur, whereas for increasing values of the selection coefficients, lower levels of phenotypic robustness are required (Fig. 1). *ρ_c_* can vary from –∞ to 1. When *ρ_c_* ≤ 0, no robustness is required for adaptation. This happens for low mutation rates and for high selection coefficients, but for whole-genome genotypes this is not a common combination (see Discussion).

**Figure 1:**
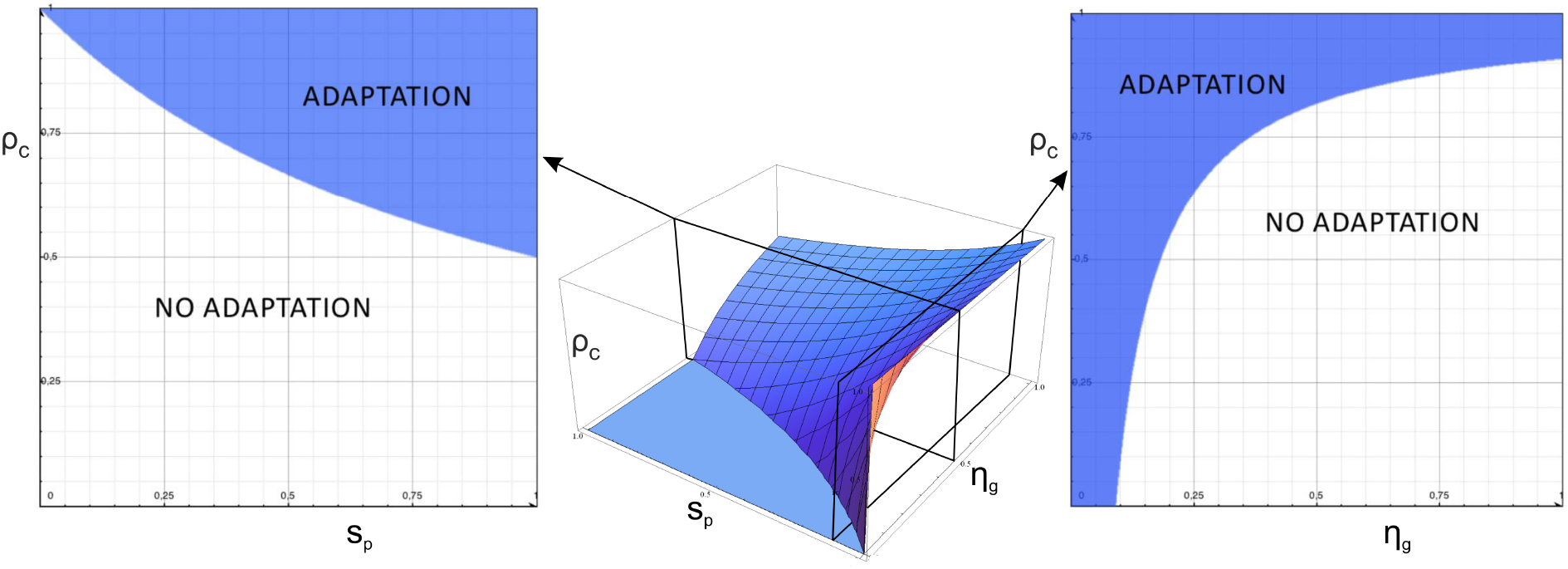
Center: three-dimensional representation of the critical robustness *ρ_c_* for different combinations of *s_p_* and *η_g_*; Left: critical robustness (boundary between empty and shaded areas) under different selection coefficients, with fixed *η_g_* = 0.5. The shaded area represents the parameter space where adaptation can occur, while the empty one where adaptation cannot occur. Right: critical robustness (boundary between empty and shaded areas) under different genome mutation probability with fixed *s_p_* = 0.1.

Studying the condition for *ρ_c_* > 0, from (11) we get

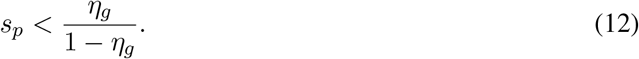

Since 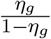 increases nearly exponentially with *η_g_* (actually, super-exponentially after 0.9), inequality (12) is often easily satisfied, and some level of robustness is required for adaptation to occur irrespective of the selection coefficient value. Moreover, as for large genomes and/or large per-base mutation rates *η_g_* tends rapidly to 1, the condition for adaptation to occur can be approximated to

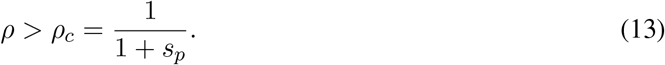

This means that the phenotypic robustness needed for a particular advantageous phenotype to spread throughout the population is inversely related to its selective advantage (*s_p_*) in that particular moment.

## 5 Discussion

Previous studies remarked the role of phenotypic robustness in enhancing evolutionary adaptation through the effect of cryptic genetic variation (e.g., Hayden et al., 2011; Rigato and Fusco, 2016), in particular as long-term effects on on evolvability (Payne and Wagner, 2019). However, on a short-term scale, i.e. on a time scale of a few generations (see Walsh and Lynch, 2018), phenotypic robustness is thought to oppose the process of adaptation by buffering the effects on favourable mutations. Here we have shown that, counterintuitively, not only phenotypic robustness can boost the adaptation process, but that it can also be a necessary condition for adaptation to occur and to be maintained in the course of evolution. There is a critical level of phenotypic robustness below which evolutionary adaptation cannot regularly occur, even in the case of sizably selection coefficients and in virtually infinite-size populations, as this treshold does not depend on genetic random drift. Critical robustness is analogous to the so called *error threshold* of the ordinary quasi-species model (Eigen et al., 1989; Wilke, 2005; Nowak, 2006), a deterministic effect of high mutation rates which impedes populations to reach and/or reside on a fitness-landscape peak, and disperse them over the sequence space. Critical robustness differs from the latter for considering genotypes and phenotypes as two distinct (although connected) spaces of organismal variation.

In Reidys et al. (2001) and Takeuchi et al. (2005) a phenotypic error threshold was discussed with respect to a parameter, λ, which represents the fraction of neutral mutants of a focal phenotype, i.e. the fraction of genotype neutral neighbours for a given genotype. Reidys et al. (2001) showed that a rather low degree of mutational neutrality can increase the error threshold unlimitedly, whereas Takeuchi et al. (2005), by considering different parameter values with respect to Reidys et al.’s model (e.g., the probability of correct replication per base), showed that the increase of the error threshold due to mutational neutrality is limited. However, although both contributions focus on evolutionary dynamics at the phenotypic level, these maintain the implicit assumptions of the original (genotype-based) quasispecies model, i.e. relatively short sequences (like those of RNA molecules and virus genomes), high selection coefficients and a single-peak fitness landscape. Here, these assumptions were relaxed by adopting a definition of robustness which does not coincides with λ but for special cases (e.g., that of a single point mutation per generation in the whole genotypic sequence), and a definition of adaptation that does not depend on equilibrium analysis. Robustness, as defined here, simply stems from considering genotypes and phenotypes as two distinct (although connected) spaces of organismal variation, with no need to further modeling either mutation patters or epistasis. This is therefore more suitable for discussing the role of robustness at the organismal level in the whole tree of life.

Critical robustness results to be directly dependent on mutation probability and inversely dependent on selection coefficient. This relationships, in combinations with the observed values of these factors in many organisms, concurs to the fact that in most biological cases, a sizable level of robustness is required. On the basis of the operational definition of genotype adopted here, where the genotype includes the whole genome of the organism (*omigenic model*; Boyle et al., 2017), the relevant mutation rate is that of the whole genome per generation. These values obviously tend to be sizably higher than the mutation rate of single genes. In multicellular eukaryotes this in the order of several mutations per genome per generation (Drake et al., 1998). As for the selection coefficient, this can vary widely, depending on the taxon, population, season, life stage, and many other factors. However, it is widely accepted that selection coefficients tend to be relatively small in nature (Orr, 2005). For example, experimental measurements of *s* usually span between 10^−4^ and 10^−1^(Tamuri et al., 2012; Nielsen and Yang, 2003; Mathieson and McVean, 2013). Small selection coefficients are also generally assumed in most evolutionary models (e.g., Tachida, 2000; Kingsolver et al., 2001; Wild and Traulsen, 2007; Wu et al., 2010). Using representative real data on the mutation rates per base pair per generation (*μ*; Drake et al. 1998; Denver et al. 2004) and genome size (*G*; Drake et al., 1998), one can easily get a rough estimation of the mutation probability per genome per generation (*η_g_*) under standard Binomial distribution of point mutations as *η_g_* = 1 – (1 – *μ*)^*G*^. Considering a selection coefficient of *s_p_* = 0.1, which represents a large, challenging value for our model, we can calculate *ρ_c_* for different kinds of organisms using equation (11). *ρ_c_* values are typically high for RNA viruses (*ρ_c_* = 0.89; *G* = 10^4^; *μ* = 10^−4^) and pluricellular eucaryotes (*C. elegans, D. melanogaster, M. musculus, H. sapiens*; *ρ_c_* = 0.86 to 0.91; *G* = 10^8^ to 10^10^; *μ* = 10^−8^), but result to be negative for DNA viruses (*ρ_c_* = –23; *G* = 8 × 10^4^; *μ* = 5 × 10^−8^) and both prokaryote and eukaryote unicellulars (*E. coli, S. cerevisie, N. crassa; ρ_c_* = –35 to –30; G = 5 × 10^6^ to 4 × 10^7^; *μ* = 7 × 10^−11^ to 2 × 10^−10^). As we have defined robustness as a probability, *ρ_c_* values <= 0 indicate that no robustness is required in the latter cases. However, for smaller and more common selection coefficients (*s_p_* < 0.01) or in consideration of the fact that during a stressful condition (and thus adaptation) virus and unicellulars can experience augmented mutation rates (from three to ten-fold the basal; Drake et al., 1998; Galhardo et al., 2007; Foster, 2007), *ρ_c_* values tends to get positive in all cases. For instance, for a bacterium like *E. coli*, in a stressful condition with a ten-fold mutation rate (*μ* = 5 × 10^−9^), and a still large selection coefficient of *s_p_* = 0.01, the minimum level of robustness required is *ρ_c_* = 0.60.

These results, which represent an attempt to formally include phenotypic robustness in the more inclusive framework of adaptive dynamics, are in agreement with the pervasiveness of robustness (often also called *redundancy*) at different levels of biological organization, from molecules to whole organisms (e.g., Rennell et al., 1991; Edwards and Palsson, 2000; Sinha and Nussinov, 2001; Giaever et al., 2002; Raser and O’shea, 2005; Raj et al., 2006; White et al., 2013; Vachias et al., 2014; Fares, 2015; Klingenberg, 2019). Phenotypic robustness qualifies as a key feature of the organism genotype-phenotype map, a major quantitative determinant of biological system’s ability to adapt and, in the end to evolve.

## Aknowledgements

This work has been supported by a grant from the Italian Ministry of Education, University and Research (MIUR) to GF.

